# A hyperthermic seizure unleashes a surge of spreading depolarizations in *Scn1a* deficient mice

**DOI:** 10.1101/2022.10.10.511466

**Authors:** Isamu Aiba, Yao Ning, Jeffrey L. Noebels

## Abstract

Spreading depolarization (SD) is a massive wave of cellular depolarization that slowly migrates across the brain gray matter. Cortical SD is frequently generated following brain injury, while less is understood about its potential contribution to genetic disorders of hyperexcitability, such as *SCN1A* deficient epilepsy in which febrile seizure often contributes to disease initiation. Here we report that spontaneous SD waves are a predominant EEG abnormality in the *Scn1a* deficient mouse (Scn1a^+/R1407X^) and undergo sustained intensification following a single hyperthermic seizure. Chronic DC-band EEG recording detected spontaneous SDs, seizures, and seizure-SD complexes in Scn1a^+/R1407X^ mice but not wild-type littermates. The SD events were infrequent, while a single hyperthermia-induced seizure robustly increased SD frequency over four-fold during the initial postictal week. This prolonged neurological aftermath could be suppressed by memantine administration. Video, electromyogram (EMG), and EEG spectral analysis revealed distinct neurobehavioral patterns; individual seizures were associated with increased motor activities, while SDs were generally associated with immobility. We also identified a stereotypic SD prodrome, detectable over a minute before the onset of the DC potential shift, characterized by increased motor activity and bilateral EEG frequency changes. Our study suggests that cortical SD is a pathological manifestation in *SCN1A* deficient epileptic encephalopathy.

## Introduction

Spreading depolarization (SD) is a massive self-regenerative wave of sustained near-complete cellular depolarization slowly propagating across brain gray matter (1, 2). The profound cellular depolarization leaves prolonged hypoperfusion and depression of spontaneous neuronal activity, contributing to neurological dysfunction of variable severity depending on genetic, humoral, and physiological predispositions. SD is frequently generated in patients after brain injury and is associated with acute and chronic neurological deficits (2, 3). SD is also associated with visual and motor deficits in some genetic migraine with aura syndromes (4), and experimental studies suggest that repetitively provoked SD in healthy brain can produce migraine comorbidities such as photosensitivity, hyperalgesia, and anxiety (5, 6). There is a growing interest in understanding SD generation mechanisms, their neuropathological consequences, and therapeutic interventions.

Due to their excitatory nature, SD events are detected in association with seizures or epileptic discharges in both experimental and clinical settings (7, 8). These “peri-ictal” SD events have been considered as a candidate mechanism contributing to epilepsy comorbidities such as peri-ictal migraine headache and immobility (9), and the appearance of SD in subcortical structures correlates with cardiovascular and respiratory dysfunction linked to postictal mortality in experimental animal studies (10–12). SD generation patterns vary depending on the pathological context. In acutely injured brains, isolated seizures, SD, as well as seizure-SD complexes coexist in some patients, while some show only a seizure or SD pattern, suggesting independent thresholds for these depolarizing events (13). Whereas recurrent spontaneous seizures define epilepsy, the characterization of SD incidence in epilepsy, especially in cases without physical injuries, is extremely limited, since the DC potential shift is difficult to detect with human scalp EEG recording of filtered signals. The advance of defined monogenic mouse models of epilepsy offers the opportunity to circumvent this limitation.

Mutations in ion channel genes are a common cause of developmental epileptic encephalopathy (DEE) (14, 15), and both gain- and loss-of-function mutations in the *SCN1A*/Nav1.1 gene are frequently identified across a range of DEE phenotypes (16). *Scn1a*/Nav1.1 channels are widely expressed in both human and rodent neocortices (17–19). However, their dysfunction preferentially impacts specific neuronal populations, such as fast-spiking inhibitory neurons due to their predominant reliance on *Scn1a*/Nav1.1 containing channels (20–22). Gain-of-function *SCNA1* mutations have been identified in rare genetic DEE and hemiplegic migraine type-3 (FHM3) patients (23, 24), and increased SD susceptibility has been reported in the knock-in mouse models (25, 26). The more common *SCN1A* loss-of-function mutations are associated with epileptic encephalopathies (e.g. Dravet syndrome) with a range of severity partly reflecting the severity of mutation (27). In these cases, reduced inhibitory neuron excitability results in network synaptic disinhibition and the emergence of hyperexcitable neuronal circuitry (20) and high susceptibility to hyperthermia dependent seizures that are believed to initiate and exacerbate epileptic encephalopathy (28). It is not known whether cortical SD is involved in the pathophysiological phenotype of this loss-of-function sodium channelopathy.

In this study, we identified the spontaneous incidence of SD in adult *Scn1a*^+/R1407X^ (hereafter Scn1a^+/RX^) mice using DC-band cortical EEG recording. During prolonged monitoring studies, we found that spontaneous SD are profoundly increased in the aftermath of a single hyperthermic seizure, and the effect was partially mimicked by a single subconvulsive pentylentetrazole (PTZ) stimulation. The exacerbation persisted for up to a week and was mitigated by concurrent administration of memantine. We also established that these SD events are not purely electrographic but affect spontaneous behaviors; cortical detection of SD is preceded by a brief period of motor hyperactivity accompanied by a high-frequency shift of EEG activity and followed by behavioral arrest and EEG suppression. Together, our study reveals a pathophysiological phenotype of *SCN1A* deficient encephalopathy and suggests SD as a potential and targetable contributor to co-morbidity mechanisms in this disorder.

## Results

### Chronic monitoring of spontaneous cortical SD and seizures in *Scn1a* deficient mice

We characterized SD incidence in *Scn1a* deficient (Scn1a^+/RX^) and WT mice using a DC-band chronic EEG recording method reported in our previous study (8). In the study cohorts, we recorded a total of seventeen Scn1a^+/RX^ (7 males, 10 females) and eleven littermate WT mice (5 males and 6 females) from a total of 13 litters. Recordings started at the age of P40-96. After 7 days of baseline recordings, mice were subjected to a single hyperthermic seizure (detail described later), and the effect was monitored for 10 days.

The results are summarized in **Figure 1**. We detected three abnormal baseline spontaneous events: isolated SD, seizures, and seizures with postictal SD complexes (hereafter “seizure+SD”) in which an SD emerged within minutes following a seizure (**Figure 1A-C**). During the study, one Scn1a^+/RX^ mouse died during hyperthermic seizure, and six mice died due to postictal sudden death or moribund condition (**Figure 1D**). While the baseline incidence of seizures and SD was relatively rare (mean of both events 0.57 ± 0.35 /day), SD was detected more commonly than seizures in most mice (**Figure 1E-F**). No seizures or SD were detected in WT mice during the baseline or post-hyperthermia periods (**Figure 1D**).

**Figure 1.**
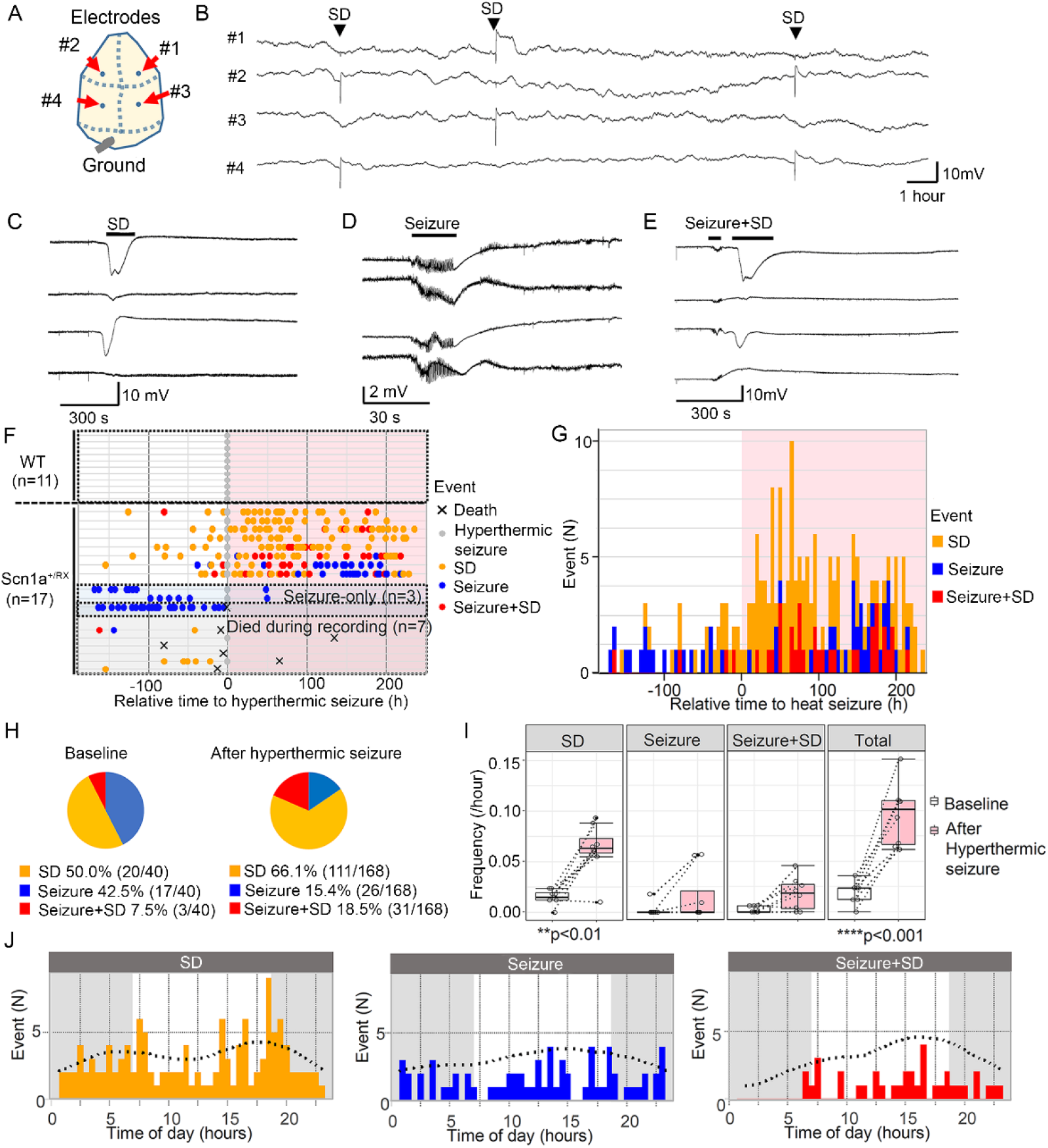
Seizure and SD phenotype of Scn1a^+/RX^ mice before and after a hyperthermic seizure. **A.** Electrode positions: from the top, #1 right anterior, #2 left anterior, #3 right posterior, #4 left posterior. **B.** Compressed trace showing a 24-hours recording. SD are reliably detected as sharp negative shift over stable baseline. **C-E** expanded representative traces of SD, seizure, and seizure+SD complex. **F-H** hyperthermic seizure robustly increased SD and seizure incidence. **F.** Raster plots of seizure and SD incidence in WT and Scn1a^+/RX^ mice. WT mice had no seizure of SD. Three mice exclusively had seizures (“seizure-only”). Seven mice died or became moribund during the study. The same Scn1a^+/RX^ even data is presented in a cumulative histogram (**G**) and pie chart (**H**) showing proportion of seizure, SD, and seizure+SD events during baseline and after a hyperthermic seizure in Scn1a^+/RX^ mice that survived the recording period, excluding the “seizure-only” mice. **I.** Quantification of event frequencies. Frequencies of SD and total events were increased after a hyperthermic seizure. “Seizure-only” mice were excluded from this analysis. Two– way ANOVA and post hoc Tukey test. **J.** Chronological analysis of SD, seizure, and seizure SD events.

After 7 days of baseline recording, mice were subjected to a hyperthermic seizure (see Methods), and its effect was continuously monitored for >10 days. Starting one day after the post-hyperthermic seizure, SD frequency clearly increased in Scn1a^+/RX^ mutants (**Figure 1D&E**), even in two mice that had not shown any seizure or SD during baseline recording. This upsurge in SD frequency lasted for multiple days and up to a week, rendering SD the dominant abnormal EEG event (**Figure 1F**). Statistical analyses detected significant increases in the frequencies of spontaneous SD (4.2-fold) and total events (3.3-fold) following the hyperthermic seizure (**Figure 1G**). In the two exceptional “seizure-only” mice that did not show SD during baseline nor during the hyperthermic seizure, seizure frequency decreased after the hyperthermic seizure. Because of the rare chance to encounter them, we could not specifically analyze more of these mice. In a time-control study in which five Scn1a^+/RX^ mice were continuously monitored without induction of hyperthermic seizure, no significant difference in seizure/SD frequency was detected (data not shown).

There was a weak diurnal trend in the SD frequency in this monitoring cohort, with peaks detected at 6.5 h and 17.8 h which correspond to light-dark cycle transitions (**Figure 1H**). A similar circadian SUDEP mortality pattern was previously detected in the same *Scn1a* deficient mouse model (29), implying the presence of a circadian vulnerability in this mutant mouse model.

### High SD susceptibility during the hyperthermic seizure in Scn1a^+/RX^ mice

In addition to its high spontaneous incidence, SD generation was also common during hyperthermic seizure in Scn1a^+/RX^ mice. Following heat exposure, Scn1a^+/RX^ mice showed cortical spikes and sialorrhea (drooling) as the body core temperature approached 40°C. Seizure onset in Scn1a^+/RX^ always coincided with a vocalization followed by a generalized tonic clonic seizure, and electrographically detected as a train of ictal spiking that resulted within less than 1 minute in a temporally coupled SD (**Figure 2A**). Mice are usually unconscious or semiconscious during the minutes-long postictal period. SD was detected in 77% (10/13) of the Scn1a^+/RX^ mice during hyperthermic seizure induction (**Figure 2C**) and was usually associated with postictal loss of voluntary motor behavior and an occasional myoclonic jerk. SD always appeared after the electrographic seizure and was likely generated as a secondary consequence of neuronal discharges, while brain hyperthermia might facilitate it. The three Scn1a^+/RX^ mice that did not display SD during hyperthermic seizure were “seizure-only” mice and one of these died shortly after this period. In WT mice, ictal SD was less common during the hyperthermic period (27% (3/11), p=0.037 vs Scn1a^+/RX^, Fisher’s exact test, **Figure 2B&C**). Consistent with earlier studies (30), the hyperthermic seizure threshold was significantly higher in WT littermates (WT: 43.0 ± 1.0°C, n=11, Scn1a^+/RX^: 41.7 ± 0.7°C, n=12, p=0.007, Mann-Whitney test) which also required longer heat exposures (>30 minutes) until a seizure emerges (**Figure 2D**).

**Figure 2.**
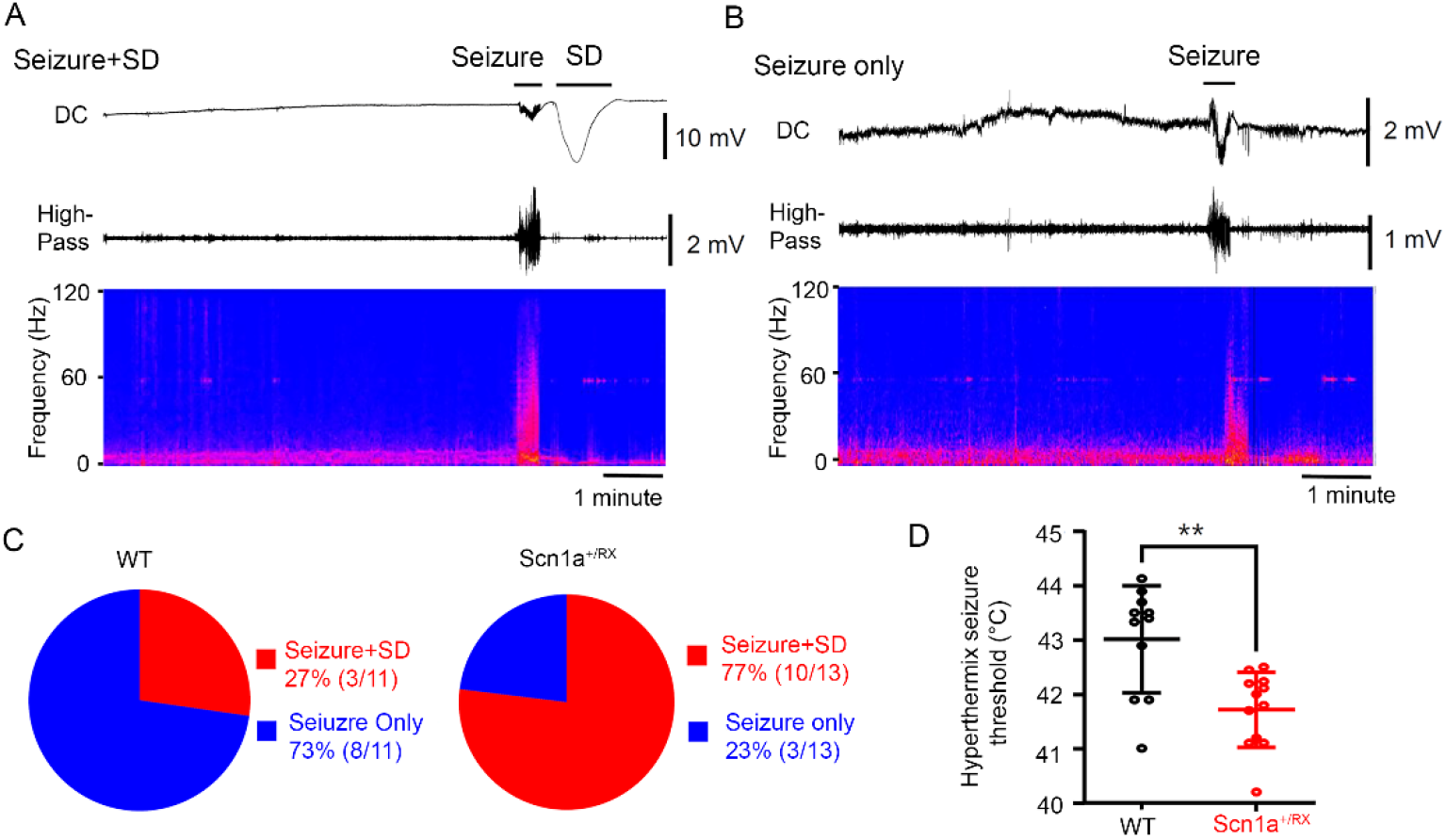
SD generation during hyperthermic seizures induced with a heating lamp in Scn1a^+/RX^ mice. **A&B** Representative EEG showing seizure and SD. *top* DC, *middle* high pass (>1 Hz), and *bottom* power spectrum of EEG (anterior electrode). **C**. Postictal SD was less common in WT mice. 82% (10/13) of Scn1a^+/RX^ mice developed SD following seizure, while 27% (3/11) in WT mice. **D.** Consistent with previous studies, Scn1a^+/RX^ mice showed a lowered thermal threshold for seizure. WT: n=11, Scn1a^+/RX^: n=13, p=0.007, Mann-Whitney U-test

### Hyperthermic seizure did not modulate the kinetics of individual seizure or SD

Hyperthermia modulated the number but not the localization or kinetics of individual SD and seizure events. We detected a total of 160 SDs, 95 seizures, and 43 seizure+SD complexes. Regardless of the presence or absence of a preceding seizure, almost all SDs were serially detected first with the posterior and next with the anterior electrode (isolated SD: 99% (159/160), seizure+SD: 98% (42/43)). While all seizures were detected as bilaterally generalized events, SDs were always unilateral and tended to be detected more often in the left than the right hemisphere (SD: left 97 vs right 63, p=0.072, seizure+SD: left 26 vs right 17 p=0.39, Fisher’s exact test). These characteristics suggest the presence of a stereotypic SD-generation mechanism in Scn1a^+/RX^ mice (see Discussion).

Postictal generalized EEG suppression (PGES) is a simultaneous silencing of EEG activity across all electrodes immediately following a generalized seizure, and its presence and duration often correlate with the clinical severity of postictal deficits (31, 32). While its origin is uncertain, the loss of EEG amplitude could resemble the consequence of SD. However, in this model, PGES period could be distinguished from the SD dependent EEG depression (**Figure 3A**). In 47% (20/43) of seizure+SD complexes, we detected PGES lasting for 13.9 ± 5.0 seconds, and neither PGES incidence nor duration differ after a seizure with or without postictal SD (**Figure 3B-C**). Thus PGES in this mouse model is a separable process that did not seem to predict or correlate with postictal SD incidence. The relatively small EEG suppression after spontaneous SD in awake mice is consistent with our earlier study (8) and likely reflects the higher spontaneous brain activity compared to anesthetized preparations.

**Figure 3.**
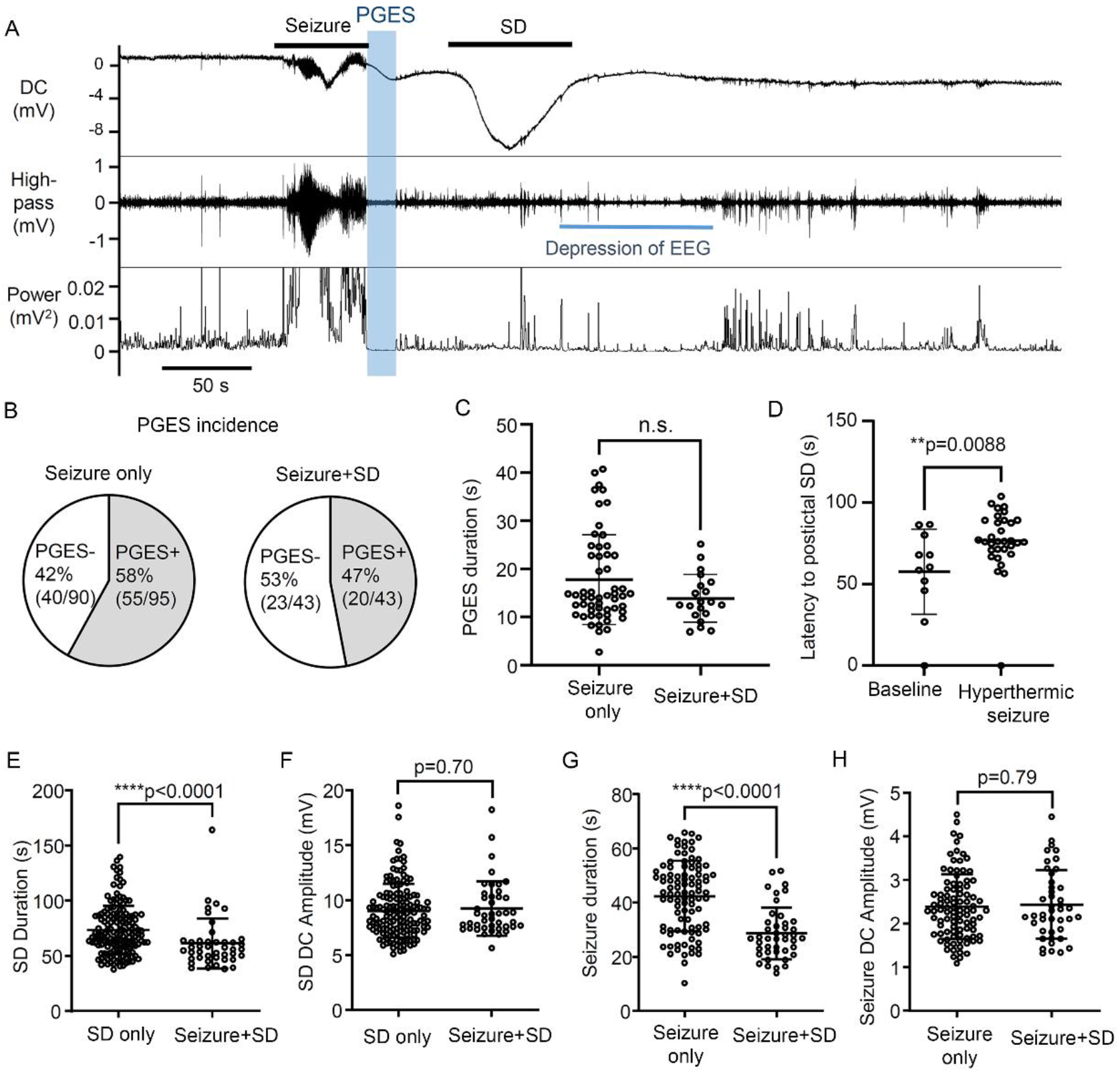
Electrophysiological characteristics of PGES and postictal SD. **A.** representative trace showing the temporal sequence of PGES (depressed EEG amplitude, blue window) and postictal SD generation. *Top*: DC, *Middle*: high pass (> 1Hz), *bottom*: EEG converted into power. The PGES incidence (**B**) and duration (**C**) were similar in seizure without postictal SD and seizure with post-ictal SD. Seizure only: n=55, Seizure+SD: n=20. **D.** The latency to SD after seizure termination is significantly prolonged after a hyperthermic seizure. Baseline: n=11, After hyperthermic seizure: n=32. **E-H** Comparison of seizure/SD kinetics between those in isolated events and those in the seizure+SD complex. The duration of SD in the seizure+SD complex is shorter than the duration of SD detected alone (**E**), while **t**he DC amplitudes were not different (**F**). Similarly, the duration of seizure in the seizure+SD complex is shorter than the duration of seizure that appeared without SD (**G**), while the DC amplitudes were not different (**H**). SD only: n=160, Seizure only: n=95, Seizure+SD: n=43. Statistics were computed by Mann-Whitney U-test

Based on the increased spontaneous SD event frequency shown in **Figure 1D**, hyperthermic seizure might be expected to facilitate postictal SD generation by shortening the time to onset. However, the latency to postictal SD was significantly prolonged after hyperthermic seizure (**Figure 3D**). Hyperthermic seizure did not alter the kinetics of individual events, thus no differences were detected in SD durations (baseline: 75.2 ± 24.7 s, n=39, after hyperthermic seizure: 72.1 ± 21.2 s, n=121, p=0.70), SD amplitudes (baseline: 9.4 ± 2.4 mV, n=39, after hyperthermic seizure: 8.9 ± 2.4 mV, n=121, p=0.18), seizure durations (baseline: 42.9 ± 13.3 s, n=44, after hyperthermic seizure: 41.9 ± 12.9 s, n=51, p=0.63), or seizure amplitudes (baseline: 2.3 ± 0.70 mV, n=44, after hyperthermic seizure: 2.5 ± 0.80 mV, n=51, p=0.46). Overall, the effect of hyperthermic seizure seems to be restricted to event generation pattern (event frequency, latency of postictal SD onset), and is without effect on the individual event severity.

In contrast, the kinetics of SD and seizure activities detected in seizure+SD complexes differ from those detected individually. The duration of SD in seizure+SD complex was significantly shorter than those SDs detected without a preceding seizure (isolated SD duration: 72.9 ± 21.8 s vs SD in “seizure+SD” 61.3 ± 22.6 s, p<0.0001, **Figure 3E**). Similarly, seizure duration was also shorter when detected in a seizure+SD complex (duration: isolated seizure 41.5 ± 13.5 s vs seizure in seizure+SD 28.6 ± 9.6 s, p<0.0001, **Figure 3G**). On the other hand, the peak amplitudes of negative DC offset during SD and seizure were unchanged (**Figure 3F&3H**).

These analyses highlight the qualitative differences between seizure and SD components in a seizure+SD complex compared to those detected individually.

### Memantine administration suppresses the prolonged aftermath of a hyperthermic seizure

We next examined whether pharmacological inhibition using the NMDAR antagonist memantine might attenuate the depolarizing aftermath of a hyperthermic seizure, since NMDAR activation has been implicated in a model of febrile seizure induced epileptogenesis (33). Memantine is an FDA-approved NMDAR antagonist, and its rapid absorption and relatively short half-life in mouse brain (∼4 hours (34)) proved useful in analyzing the temporal contribution of NMDAR to hyperthermic seizure induced SD exacerbation. We selected 10 mg/kg (i.p.), a dose that inhibits an NMDAR-dependent plasticity mechanism *in vivo* (35), and used different administration patterns to investigate critical time windows; pre-treatment (30-60 minutes before the hyperthermic seizure), post-treatment (6 and 16 hours after hyperthermic seizure induction), and a combined pre/post-treatment. The drug effect was evaluated by comparing the frequency of total events before and 24 after hyperthermic seizure induction. The initial 24-hour postictal period was excluded from the analysis to avoid the acute pro-convulsive effect of prolonged memantine administration. In fact, two mice in pre/post treatment group showed a recurrent seizure during memantine administration (**Figure 4D**, see Discussion).

**Figure 4.**
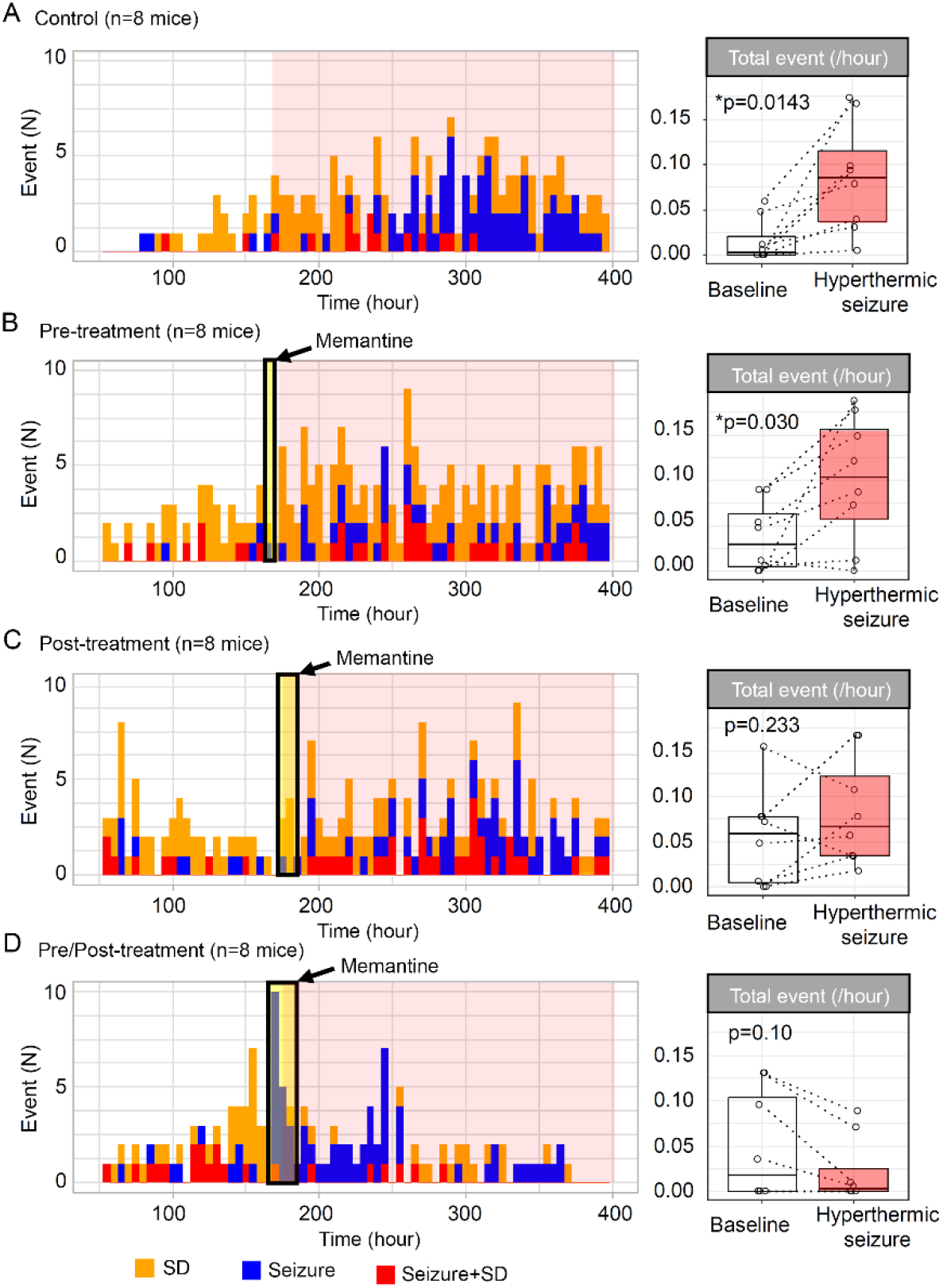
Prolonged memantine administration inhibits the hyperthermic seizure induced exacerbation of depolarizing events in Scn1a^+/RX^ mice. Cumulative histogram bars show SD incidence (orange), seizures (blue), and seizure+SD complexes (red) before and after hyperthermic seizure. Box-plots show total event frequency (total events per hour) during baseline (white) and after hyperthermic seizure (red). **A.** pattern of events in saline pretreated control Scn1a mutants. **B.** Efficacy of single dose memantine (10 mg/kg i.p.) pretreatment administered 30-60 minutes before hyperthermic seizure, **C.** efficacy of memantine post-treatment repeated 6 and 12 hours after hyperthermic seizure, and **D.** combined pre- and post-treatment data were analyzed. The duration of treatment is shown in the yellow shade, and the duration of the post-hyperthermic seizure period is in a pink shade. At right, the frequency of total events during baseline and following the hyperthermic seizure are shown. In each group, n=8 mice. Statistics calculated by paired Wilcoxon signed-rank test.

Memantine at high dosage is known to acutely inhibit SD generation in some models (36). We found that pretreatment with memantine was partially effective at inhibiting SD generation during hyperthermia seizure; SD incidence in memantine pretreated mice (i.e. pre- and pre/post memantine treated mice) was 56% (9/16 mice), which is less compared to untreated mice (untreated control + post-treatment mice, 88% (14/16 mice, p=0.052, Fisher’s exact test). Memantine administration inhibited the chronic aftermath of hyperthermic seizure (two-way ANOVA, Memantine administration paradigm: F= 1.68 p=0.18, time effect: F=5.74, p=0.020, interaction: F=2.63, p=0.060). All mice receiving saline before hyperthermic seizure showed a robust increase in the total number of events (**Figure 4A**, p=0.014). In this control cohort, more mice later developed seizures compared to the initial cohort shown in **Figure 1**. A significant increase in total events was also seen in the pre-treatment group (**Figure 4B**, p=0.030), while the aftermath was absent in one mouse. Memantine treatment initiated after hyperthermic seizure attenuated the aftermath (**Figure 4C**, p=0.23), although the total number of seizure/SD events was still increased in 75% (6/8) of mice. The combined pre/post-treatment was most effective, as the frequencies of the seizure/SD events were unchanged or decreased during post-hyperthermic seizure period in all mice tested (**Figure 4D**, p=0.10).

These results collectively suggest that continuing NMDAR activation after hyperthermic seizure contributes to the exacerbation of depolarizing phenotype of Scn1a^+/RX^ mice.

### Subconvulsive PTZ partially mimics the effect of hyperthermic seizure

The contribution of NMDAR activity raised a possibility that the prolonged increase in spontaneous SD incidence following a hyperthermic seizure might be explained by subthreshold circuit hyperexcitation. This possibility was tested by a single injection of PTZ, a convulsant that acts as a GABA-A receptor antagonist. After baseline recordings, seven Scn1a^+/RX^ mice were injected with PTZ (30 mg/kg, i.p.), a dosage considered subconvulsive in WT mice (ED_50_ ∼ 65 mg/kg (37)). Although the half-life of PTZ in mouse brain is not available, it is estimated to be <2 hours based on the analysis in dogs (38) as well as the time course of appearance of PTZ-provoked epileptic spikes in our recordings. Thus SD or seizure events appearing 2 hours after administration are considered secondary to the direct network excitation by PTZ.

Five mice showed recurrent ictal spikes without seizure or SD during the 2 hours after PTZ injection (**Figure 5A-C**), while two mice showed a seizure immediately after PTZ injection and one of them died postictally. Continuous monitoring revealed enhanced recurrent SD/seizure frequencies for several days in four surviving mice (**Figure 5B-C**) and there was a trend toward increase in the mean total seizure/SD events (**Figure 5D**, p=0.0625). This result suggests that subconvlusive synaptic disinhibition is sufficient to produce a prolonged hyperexcitation profile of enhanced SD/seizure in Scn1a^+/RX^ mice when NMDAR overactivation is present.

**Figure 5.**
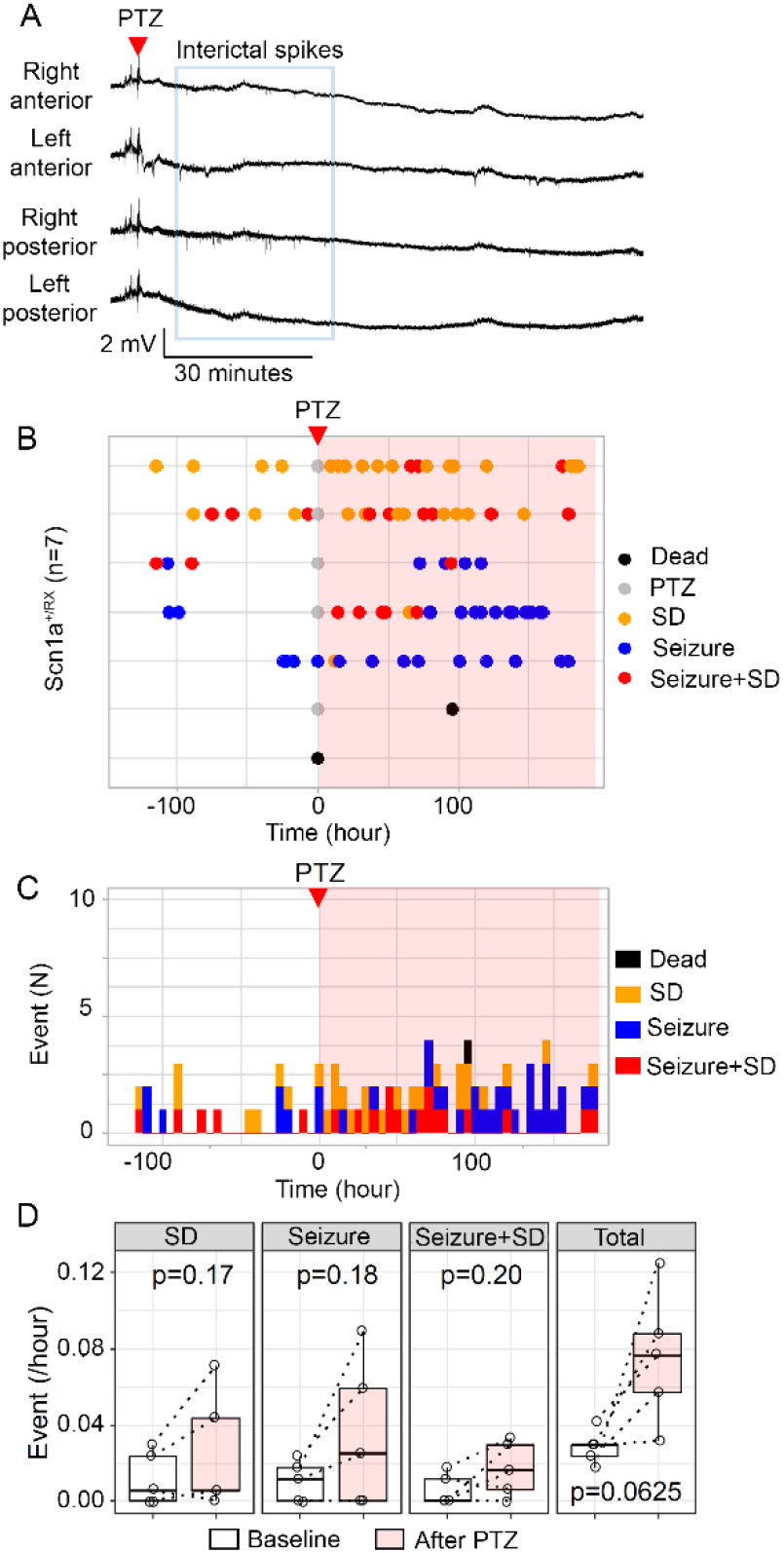
Subconvulsive PTZ stimulation partially mimics hyperthermic seizure effect. **A.** representative traces of EEG activity following PTZ injection (30 mg/kg i.p.) in seven Scn1a^+/RX^ mouse. PTZ increased interictal epileptic discharges for 30 minutes without seizure in this mouse. Traces from top: right anterior, left anterior, right posterior, left posterior. **B.** Raster plots of SD, seizure, and seizure+SD complex during baseline and after PTZ injection (pink shade). Two mice died during the recording. **C.** Same results presented in cumulative histogram of SD, seizure, and seizure+SD incidences. **D.** quantitative comparison of frequencies of SD, seizure, seizure+SD, and total events before and after PTZ injection. n=5, Statistics calculated by paired Wilcoxon signed-rank test.

### Interictal and postictal motor activity associated with SD

We next sought to identify neurophysiological deficits associated with SD using chronic EEG with neck-muscle electromyography (EMG). This analysis revealed stereotypical motor activity patterns accompanying interictal SD episodes, characterized by prodromal minutes-lasting motor activity prior to SD detection at the parietal electrode, followed by suppression of activity for minutes after SD was detected at the frontal electrode (**Figure 6A**). In contrast, seizures were associated with a robust EMG signal increase following the onset of EEG discharges (**Figure 6B**). In some mice, these convulsive EMG signals were transiently suppressed during PGES as reported in electrically evoked seizure model (39), but were then followed by bursts of motor activity. During a seizure+SD complex, the seizure again triggered an initial sharp EMG activity burst, followed by a transient reduction once SD is detected at the frontal electrode (**Figure 6C**). In the majority of mice, EMG activity remained low after the postictal SD. Overall, seizure+SD showed lower postictal motor activity compared with seizure (p<0.001, analysis of variance of aligned rank transformed data).

**Figure 6.**
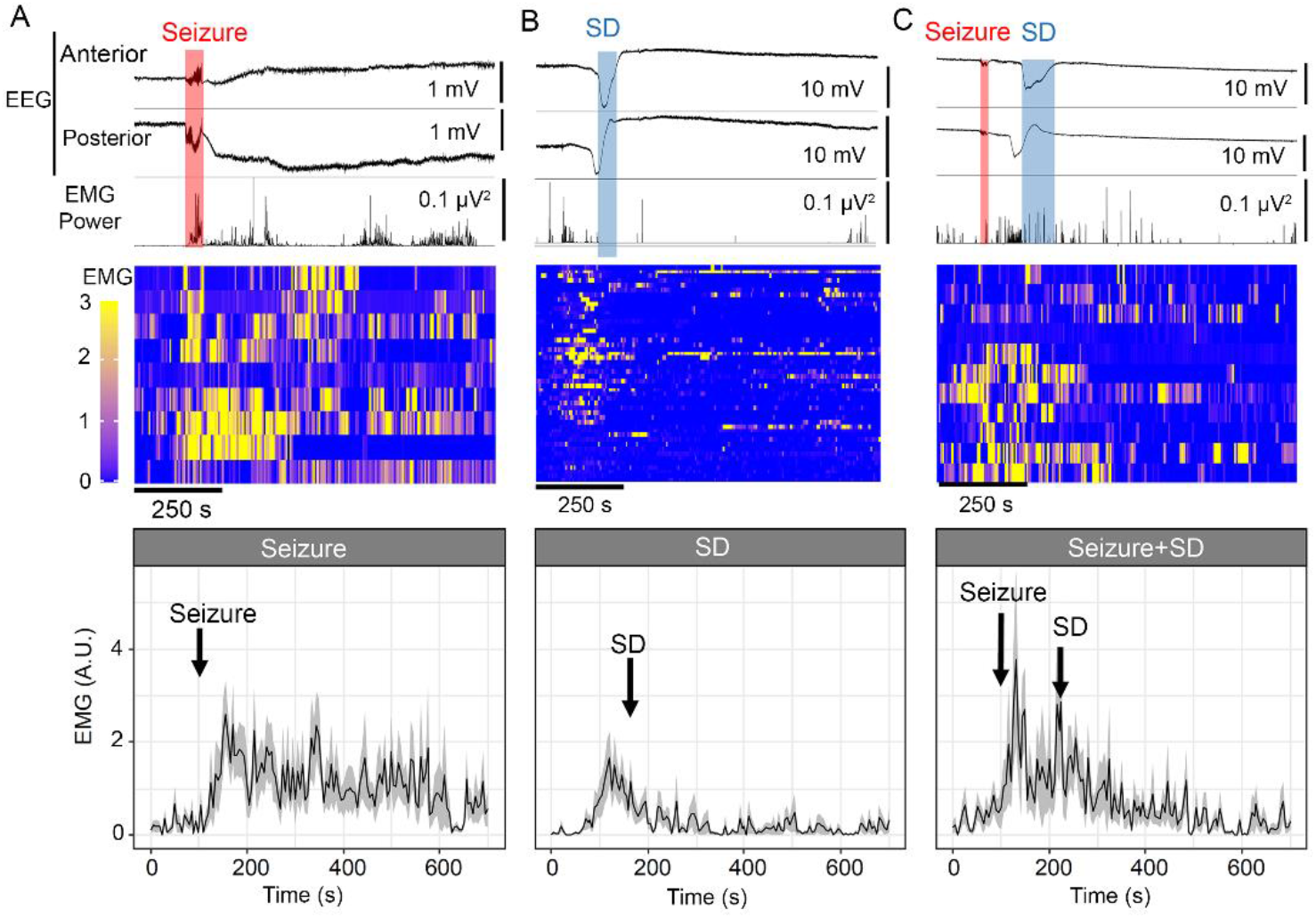
Analysis of EMG motor activity associated with SD (**A**), seizure (**B**), and seizure+SD (**C**). Top: unilateral anterior and posterior EEG and simultaneous neck EMG trace. Middle: raster plot of individual analyzed EMG signal patterns. Each lane represents a single event in a representative animal. Bottom: averaged traces of EMG activity are presented as mean ± standard error. **A.** Seizure is associated with an abrupt increase in EMG signal which is followed by motor activations. **B.** SD was associated with prodromal behavior activation, followed by suppression as DC shift is detected in the frontal cortex. **C.** Seizure+SD is also associated with initial convulsive motor activity, which is inhibited once SD is detected over the frontal cortex. Some motor activity is present after postictal SD but is reduced in comparison to seizure alone. n= 11 47, 9 events for seizure, SD, and seizure+SD, respectively. p<0.001 in EMG patterns between events, Aligned rank transformation ANOVA

### Decreased voluntary locomotor behavior is associated with SD

Locomotor behavior associated with seizure, SD, and seizure+SD events was analyzed using video recording. In total, 37 seizures, 67 SDs, and 14 seizure+SD from 7 mice were analyzed after the removal of low-quality images. Average locomotion during a 30 minutes period before and after each event type is presented in **Figure 7A**. Unlike EMG analysis (**Figure 6B**), this analysis did not detect behavior increase associated with isolated seizures, suggesting that increased ictal motor activities detected by EMG were mostly stationary jerking movements. Similar to EMG analysis, SD tended to be associated with decreases in locomotion minutes after SD, while the effect is less robust. Combined seizure+SD suppressed locomotion for a prolonged period.

**Figure 7.**
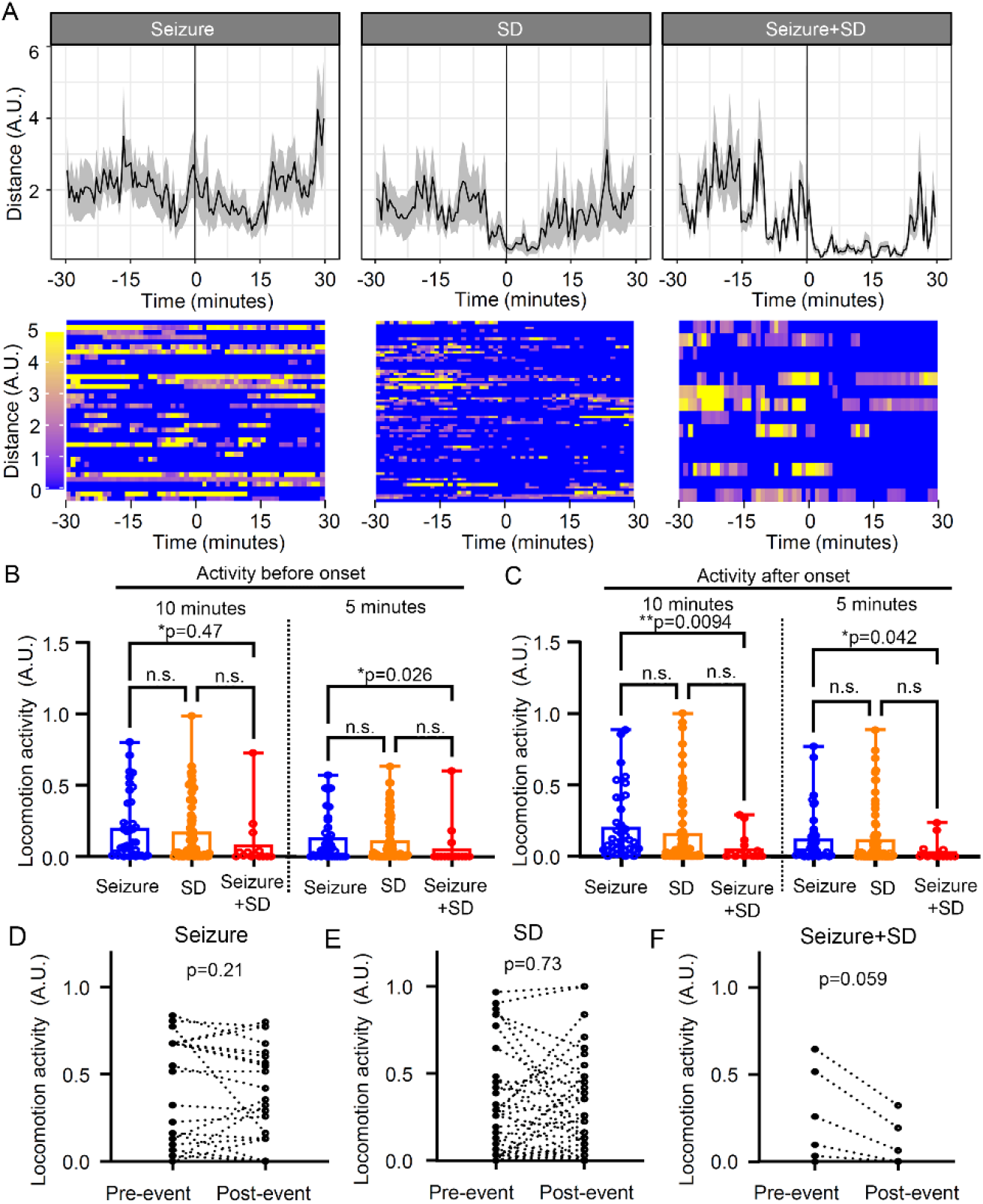
Locomotion changes associated with seizure, SD, and seizure+SD were analyzed using video images. **A.** Average traces (top, mean ± standard error) and raster plots (bottom, each lane shows a single event) 30 minutes before and after event onset (line at t=0). n= 37, 67, and 14 events for seizure, SD, and seizure+SD, respectively. **B&C.** Locomotion was also analyzed by a binary method (see Results). Comparison of locomotion 10 or 5 minutes before (**B**) and after (**C**) each event. Seizure+SD is associated with reduced pre- and post-event locomotion activity. **D-F.** Comparisons of locomotion changes in individual events 3 minutes before and after each event. Seizure and SD did not show consistent directional changes, however, seizure+SD events consistently reduced locomotion activity. Two-way ANOVA, Event: F = 2.01, p=0.14, Time: F=15.93, p<0.001, interaction: F=1.23, p=0.30.

Since the analysis of mean data could be biased by a few events with large activity, we also analyzed the locomotion using a binary method in which 5 seconds time bins are scored “active” or “inactive” periods (see Methods). The binned analysis 5 and 10 minutes before and after event onset again revealed lower total activity associated with seizure+SD compared with seizure alone, while SD alone showed an intermediate level (**Figure 7B-C**). Analysis of locomotion before and after each event did not show directional changes in seizure (p=0.21) and SD (p=0.73), while seizure+SD trended to reduce the subsequent locomotion (p=0.059, **Figure 7D-F**).

Together, these motor activity patterns suggest that SD episodes suppress voluntary motor behaviors as was previously observed in optogenetically evoked SD (40, 41).

### Stereotyped EEG activity changes also precede the detection of negative DC potential shift in the cortex

Given the behavioral evidence of a neurological precursor signaling SD onset detected with EMG analysis, we examined EEG characteristics preceding the detection of a spontaneous cortical SD using 45 randomly selected events from 10 Scn1a^+/RX^ mice. This analysis identified a striking signature preceding isolated SD episodes, characterized by a sudden and bilaterally simultaneous reduction in low frequency (0-30 Hz) and increase in high frequency (30-120 Hz) band EEG activity (hereafter termed pre-SD, **Figure 8A** p<0.0001, both SD side and contralateral hemisphere). These two frequency bands were selected based on multiple analyses to obtain robust characteristics with reproducibility and simplicity. During the pre-SD period, total EEG power (0-120 Hz) was reduced in both hemispheres as low frequency activity dominated the baseline EEG power, while the decrease was slightly larger in the SD-affected hemisphere (Pre-SD total EEG power relative to baseline: SD side: 43.9±20.0% vs Contralateral: 47.5±17%, p=0.039, n=45). This EEG power change occurred 87±18 s before the negative DC shift onset, and was temporally correlated with the prodromal motor activity increase. The onset of the pre-SD phase was always associated with compound cortical spikes (**Figure 8A** inset) detected in all cortical channels. This pre-SD EEG condition disappears once the negative DC potential shift emerges.

**Figure 8.**
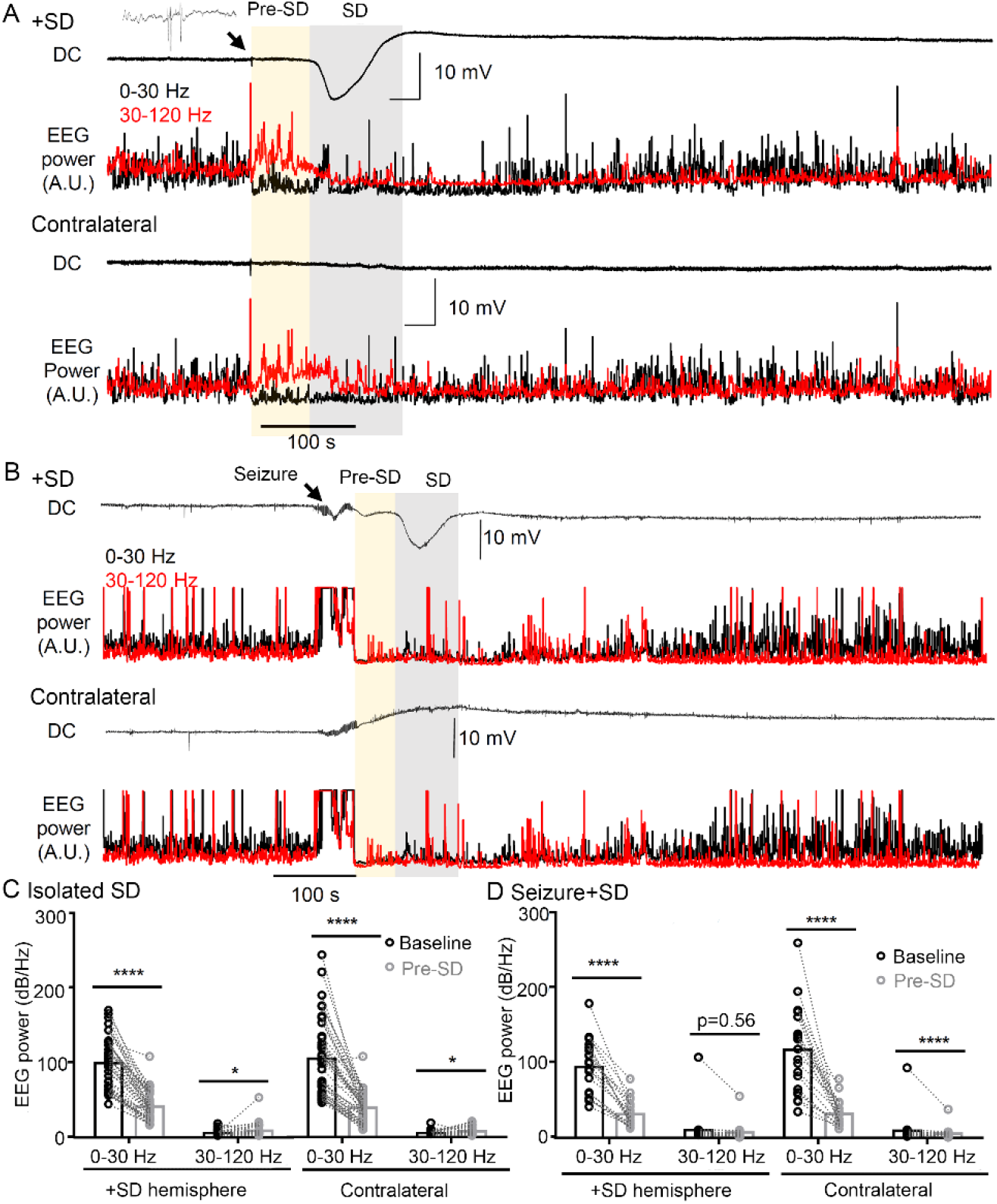
Prodromal EEG frequency change precedes the onset of the negative DC potential shift of SD. **A.** DC and EEG power changes in isolated SD. EEG activities showed a robust high frequency shift (yellow shade) more than a minute before the onset of the negative DC potential shift of SD. Note complex EEG spikes are always detected at the onset of prodromal changes. DC, Low band (0-30 Hz: in black), and high band (30-120 Hz: in red) EEG power in the SD affected (top) and the contralateral hemisphere (bottom) are shown. **B.** DC and EEG power changes during seizure+SD complex. **C.** Quantification of the EEG power during baseline and prodromal phase from 45 isolated SDs. The EEG frequency was altered in both SD affected and contralateral hemispheres (time effect: p<0.0001 both hemispheres, Repeated Measures Analysis of Variance). **D.** Same analysis of prodromal EEG frequency change in 21 seizure+SD complexes (time effect: p<0.001 both hemispheres, Repeated Measures Analysis of Variance). *p<0.05, **** p<0.001, post-hoc paired *t*-test with holm’s correction.

A similar pattern of low/high frequency change was also detected in the pre-SD phase of seizure+SD (**Figure 8B&C**, p<0.0001, both SD side and Contralateral, n=21) although high-frequency activity did not change over the SD affected side, partly because EEG activities in both low and high frequency bands were already depressed following a seizure in both hemispheres (Pre-SD total EEG power relative to baseline: SD side: 34.7±18.0% vs Contralateral: 39.1±14.7%, p=0.16).

Together, our analysis identified a clear electrobehavioral signature appearing more than a minute before cortical SD detection, likely reflecting the effect of SD generation and spread from a remote site. The EEG frequency shift and associated motor behavior changes may serve as a predictive biomarker signaling SD initiation in this model.

## Discussion

In this study, we identified a spontaneous cortical SD generation phenotype in the Scn1a^+/RX^ mouse model. SD episodes were readily detected in *Scn1a* deficient mice, usually more frequently than seizures, and the incidence of SD was robustly increased after a single hyperthermic seizure, an effect that can last for days or a week. A single subconvulsive dose of PTZ produced a similar upsurge. The post-hyperthermia week-long flurry of SD events could be inhibited by memantine when administered immediately before and continuously after the hyperthermic seizure, suggesting that early and prolonged NMDAR activation may redundantly contribute to the prolonged aftermath. Simultaneous electrophysiological and behavioral characterization also revealed that seizures and seizure+SD complexes are qualitatively different and associated with distinct prodromal and postictal behavioral states. These short-term effects may herald more consequential behavioral and cognitive deficits reported in *Scn1a* deficient mouse models (42–45). We have not analyzed younger age and cannot yet evaluate whether an early onset of SD might contribute to the developmental cognitive delay apparent in childhood Dravet syndrome. While our preclinical study suggests SD as a prominent pathological event associated with *Scn1a* deficiency, the overall clinical incidence and significance of SD in the DEE remain uncertain due to the difficulty of reliably detecting SD in human scalp EEG recordings. Further study will be required to determine whether the prodromal EEG activity we detected preceding SD initiation can potentially serve as a surrogate biomarker of SD occurrence. Alternative pharmacological SD may also help determine whether identification and early clinical management of SD will prove useful for predicting the emergence of clinical comorbidities in these individuals.

### Spontaneous cortical SD characteristics of *Scn1a* deficient mice

Our study revealed frequent spontaneous episodes of SD, rare seizure+SD complexes, and a prolonged period of high SD susceptibility following a hyperthermic seizure in the Scn1a^+/RX^ mouse cortex. Throughout the study, no SD was detected during handling or in the presence of an investigator during the chronic monitoring, thus spontaneous SD detected here appears unlinked to potential experimental stress. Acute SD generation after a heat-induced seizure was previously documented in a two-photon Ca^2+^ imaging study of immobilized Scn1a^+/-^ mice (46), and our study extended the finding by demonstrating and analyzing in detail these spontaneous events and associated behavior comorbidities. Hyperthermia facilitates SD generation in brain injured patients (47), and bath temperature above 38°C can trigger SD in isolated healthy hippocampus slices (48). This further underscores the higher SD susceptibility of *Scn1a* deficient brain, since SD during hyperthermic seizure was more common in Scn1a^+/RX^ than WT mice despite the fact that their seizures were provoked at a lower body temperature than WT.

The neurophysiological mechanism underlying the high spontaneous SD susceptibility in Scn1a^+/RX^ mice remains unclear and the exact molecular pathogenesis is likely to be a gene specific. In a model of type-2 familial hemiplegic migraine, spontaneous SD arises as a consequence of localized glutamate excess associated with genetically impaired astrocytes (49). This mechanism is unlikely to explain the pathogenesis of SD in the *Scn1a* deficient mouse where impaired GABAergic inhibition is the major mechanism of circuit hyperexcitation. We also observed that a network disinhibition by subconvulsive does of PTZ facilitated SD generation. In our study, SD was almost always sequentially detected with posterior to anterior electrodes, suggesting a stereotypic SD generation specific to the *Scn1a* deficient network disinhibition. In this regard characterization of SD phenotype in related epilepsy mutant mice with genetically disinhibited neural circuits *(e.g. GABRG2* mutations (50)) could provide further insight. Because of the low spatial resolution of our 2-electrode DC recording, we could estimate the speed of propagation of SD to be the typical range (isolated SD: 2.19 ± 0.87 mm/minute, SD in seizure+SD 2.27 ± 0.83 mm/minute), but not the propagation pattern of the SD wave. Imaging studies or a multi-electrode array may be needed to resolve this detail.

Our study detected SD events that outnumber seizures in the adult *Scn1a* deficient mouse model and most of these SD events appeared in succession without a concurrent seizure. An earlier study suggested that SD may act as an anti-seizure mechanism (51) was based mostly on the observation that SD transiently interrupts pharmacologically provoked ongoing EEG seizure discharges. We consider the mechanisms studied in that study are more relevant to status epilepticus, rather than the spontaneous SD and seizure occurrence studied in this genetic epilepsy model which are detected only a few times per day. Instead, we consider that the cortical activity state with SD and seizure represent a distinct disease profile as suggested by the consequence of hyperthermic seizure.

The frequent spontaneous SD waves detected in LOF *Scn1a* mutant mice described here may be counterintuitive, as it suggests a shared, rather than opposite SD phenotype between LOF and GOF mutant models. Our result rather suggests that SD might be commonly triggered in hyperexcitable epileptic brain even without a variant-specific mechanism as SD can be readily triggered by external stimulation in the WT mouse cortex, while some mutations would certainly facilitate SD generation and create a unique phenotype as was seen in the *Kcnq2*-cKO mice (8). Similarly, the frequent SD in Scn1a^+/RX^ mice seems contradictory to the observation that migraine with aura, generally considered to be an SD-linked neurological symptom, is not common in Dravet syndrome patients. Several possible explanations can be considered, including patient age (only 1-3% of 7 old children experience migraine, and the occurrence of aura without headache (52)). The dissociation of these phenotypes may have several explanations: The pain threshold may be higher in Dravet syndrome due to a deficiency of peripheral *Scn1a*/Nav1.1 channels maintaining sensory nerve excitation (53), whereas FHM3-linked *Scn1a* GOF mutations might reduce this threshold (54) and intensify headache sensation. FHM may represent a rare form of migraine linked to high SD susceptibility involving motor-related cortical regions, and SD generated in epilepsy may accompany distinct focal functional impairment depending on the affected brain region (55).

### EEG signatures preceding spontaneous SD in *Scn1a* deficient mice

We identified a prodromal high frequency shift in EEG activity more than a minute before emergence of the negative DC potential shift in this model. Similar high frequency activity preceding SD onset has been reported in SD evoked by potassium injection in anesthetized rat hippocampus (56). However, in that study, changes were detected only 5-10 seconds before onset of the negative DC potential shift and may not be directly comparable with the prodromal changes detected in this study. However, as in that study, we also identified complex EEG spikes at the onset of the pre-SD EEG pattern (**Figure 8A** inset) which may possibly reflect a triggering mechanism for SD in this model.

In our recordings, more than 97% of spontaneous SD events were detected sequentially from the posterior to anterior electrodes, indicating a stereotypic SD generation pathway in this mouse. The result seems inconsistent with the known high anatomical SD susceptibility zone in the somatosensory barrel cortex in rodents (57, 58) which is located closer to the anterior electrodes. This discrepancy might indicate that spontaneous SD generation in Scn1a^+/RX^ mice differs from the previous experimental SD models. Due to the limited spatial resolution in our chronic monitoring study, we could not pinpoint the spontaneous SD generation site more precisely. A microcircuit-based study of this mouse model may help identify an SD hotspot and better define the SD generation mechanism.

Our study also found that the duration of seizure activity during a “seizure+SD” complex is shorter than those in isolated seizure events. Given the stereotypic prodromal changes we found, the emergence of postictal SD may be somehow pre-determined by this altered physiological state. We also identified a stereotyped delay in postictal SD onset, almost always one minute after seizure termination. This latency is in contrast with the *Kcnq2* conditional KO mouse model, where most SDs were bilateral and generated even before seizure activity fully terminated (8). This result highlights the distinct SD regulatory mechanisms between epilepsy models with specific ion channel defects.

### Aggravation of SD phenotype by a hyperthermic seizure

Febrile seizure due to infection or prolonged heat exposure is a common mechanism contributing to the onset and progression of DS symptoms, and similar hyperthermia-induced disease exacerbation is recapitulated in juvenile *Scn1a* deficient mice (59, 60). Our observation is generally consistent with these earlier studies and further extend them by demonstrating that 1) a single brief seizure is sufficient to provoke an upsurge of SD, 2) seizure and SD exacerbation can occur in adult mice, and 3) the hyperthermic seizure effect can be prevented by concomitant memantine administration. This evidence suggests that the pathogenic mechanism of disease progression by a hyperthermic seizure may occur independently of critical developmental stages. In other words, the evolution of *Scn1a* deficient encephalopathy may reflect sustained proexcitatory plasticity mechanisms rather than being solely determined by a static developmental defect. This view is consistent with studies demonstrating that postnatal conditional *Scn1a* deletion can create an epileptic encephalopathy similar to the genomic model (61, 62), and also that the disease phenotype can be reversed or ameliorated when *Scn1a* expression is recovered at a juvenile age (43, 63, 64).

The sustained proexcitatory effect of a hyperthermia-provoked seizure compared to the less consequential spontaneous seizure suggests an underlying functional difference between endogenous and exogenously triggered pathological excitation. In addition, two seizure-prone mice that had a reduced seizure frequency after hyperthermic seizure (**Figure 1**) suggest the effect of hyperthermic seizure could be altered in distinct disease conditions. Although the number is limited due to the rare chance to encounter these mice, they showed no SD during chronic monitoring and hyperthermic seizure. In addition to having seizures, increased SD threshold might correlate with the altered response to hyperthermic seizure.

### Behavioral consequences of spontaneous SD

Our study also revealed that spontaneous SD is associated with motor behavior changes. SD was preceded by prodromal activation and this motor behavior is later suppressed when SD invades the frontal cortex (**Figure 5**). The latter is generally consistent with findings from optogenetically-evoked SD in awake animals (40, 41). The seizure+SD complex also showed prolonged motor behavior suppression which was occasionally detected as continuous immobility following PGES. A study in α2-Na^+^/K^+^ ATPase conditional KO mice described a paralyzing effect of putative cortical SD events (65). However, the paralytic state described in that study more closely resembled postictal coma than immobility. Postictal immobility has also been linked to the risk of severe respiratory dysfunction and sudden unexpected death in epilepsy (SUDEP)(31), a comorbidity in Dravet syndrome cases and mouse models. It will be important to examine whether the presence of postictal SD also modifies the cardiorespiratory consequences of seizure. Optogenetically evoked cortical SD can modify the heart rhythm in a sleep/awake status-dependent manner (40) and our unpublished observation).

Beyond the acute behavioral effect identified here, the early developmental and lasting effects of SD on the global neurological disorder remain to be elucidated. Previous studies suggest recurrently provoked SD can injure synaptic biology, and incur chronic deficits associated with migraine with aura, such as hyperalgesia, photophobia, and anxiety (5, 6). In the current study design, we could not accurately measure these parameters because such behavior tests interrupt the EEG recording and potentially affect the seizure/SD generation pattern. More controlled behavioral characterizations will be needed to understand the long-term contribution of spontaneous SD to neurological deficits associated with this developmental epileptic encephalopathy.

### Benefits and limitations of memantine treatment

The demonstration that memantine can inhibit the hyperthermic seizure response (**Figure 4**) suggests a contribution of sustained NMDAR activation to the prolonged aftermath. Maximal suppression was found when memantine was administered before the hyperthermia episode. This effect is consistent with a recent randomized placebo-controlled clinical trial in which chronic memantine treatment reduced seizure frequency in pediatric DEE patients (66). The prolonged suppression of SD/seizure frequency by memantine administration seen in our study suggests memantine may have a potential role in DEE therapy when carefully administered.

On the other hand, we noted several *Scn1a* deficient mice that displayed recurrent seizures activity during memantine administration, which resolved after the treatment. This response is probably related to reports of a paradoxical convulsive or excitatory effect of NMDAR inhibition. For example, acute NMDAR inhibition preferentially inhibits GABAergic interneurons (67), while chronic inhibition or depletion increases neuronal synaptic and intrinsic excitability (68–72), and some loss of function NMDAR mutations are associated with developmental epilepsy (73). Determining the lowest effective dose and slow escalation of the memantine dose may circumvent these events. A better understanding of the underlying cellular and molecular mechanisms linked to postictal sustained NMDAR activation, such as inflammation (74, 75) and neurotrophic signaling (76), may identify additional treatment options.

The present study identified a prominent SD phenotype and disturbed motor behavior associated with these depolarizing events in the *Scn1a* deficient mouse model of Dravet syndrome. Together with our recent study of potassium channel deficient mouse models of DEE (8), SD may represent a neurological mechanism underlying the complex neurological deficits associated with genetic epileptic encephalopathy.

## Materials and Methods

### Animals

The Scn1a^+/R1407X^ mice, originally developed in Dr. Yamakawa’s laboratory (21), were backcrossed with the heterozygote flox-tdtomato line (100% C57BL6J background, Ai9, Jackson Laboratory #007909) and maintained on 87.5% C57BL6J and 12.5% SV129 background. The flox-tdtomato allele is inactive in the absence of Cre recombinase and was used as a post-hoc label to blind the experimenter to genotype or treatment. Littermate wild-type (WT) mice were used as controls. During breeding and the chronic recording study, the mice have ad-lib access to mouse chow (5V5M, PicoLab) and drinking water in rooms maintained at 20-24°C and 30-50% humidity with a 12-hour light-dark cycle.

### Surgery

Mice were injected with meloxicam (5 mg/kg, s.c.) as a presurgical analgesic, and anesthetized with isoflurane (3.0% induction, 1.5-2.5% maintenance) while body temperature was maintained with a heating mat. The scalp hair is depilated and the skin cleansed with betadine and 70% ethanol 3 times. After injection of local anesthetics (1% lidocaine, 1% bupivacaine mixture), a midline skin incision is made to expose the skull surface, connective tissues are removed with cotton swabs. Cranial burr holes (∼1 mm diameter) were made 0.5 mm anterior/2.0 mm lateral and 2.5 mm posterior/2.5 mm lateral from the bregma and two burr holes in the occipital bone; the position occasionally required adjustment by ∼0.5 mm when major blood vessels were present. In each burr hole, a drop of dexamethasone (10 mg/ml) was topically applied to ameliorate tissue inflammation. Teflon-insulated silver wires (36-G or ∼0.13 mm diameter) were carefully positioned over the cortical surface and a pair of wires were inserted under the neck muscles for electromyogram (EMG) recording. After implantation, the wires and connector are cemented to the skull with Metabond. Mice received meloxicam (5 mg/kg) for 3 postoperative days. To minimize the postsurgical effect, EEG recordings were initiated at least 7 days after the implant.

### Video EEG recording of awake mice

EEG recordings were conducted in an IACUC-approved satellite room (20-22°C, 40% humidity, 12-hour light-dark cycle). Each mouse was connected to an EEG tether wire and housed in a 30 x 15 cm home cage with freely available water and food. To minimize the potential influence of exposure to a new environment, activity recorded during the first 48 hours in the cage was excluded from the analysis. EEG activity was acquired using a Bioamp DC amplifier at 1 kHz and digitized by the LabChart system (ADI) while the behavior was continuously recorded using an IR-light equipped CMOS camera at 2 Hz. EEG signals were analyzed using LabChart (ADI) and Clampfit 10 software (Molecular devices). The use of DC-compatible amplifiers was necessary for SD detection by cortical surface EEG recording as slow signals are filtered out in most AC-coupled amplifiers.

All raw EEG traces were visually inspected for seizure and SD incidence and were manually quantified based on their characteristic EEG patterns as described in our previous study (8). Postictal generalized EEG suppression (PGES) was defined as the complete suppression of EEG amplitude immediately after seizure termination and the duration was determined based on the total EEG power (total EEG power <0.01 mV^2^, see **Figure 3A**). In the EEG analysis in **Figure 8**, the total power of low-frequency (0-30 Hz) and high-frequency band (30-120 Hz) cortical activities were calculated by fast Fourier transformation (N=1024, cosine-bell waveform).

The video image was analyzed using Bonsai and R software. Locomotion was calculated based on the horizontal displacement of the mouse body centroid. Some video images were excluded from the analysis because the mouse position could not be accurately resolved. The peaks of EEG event incidences in the circadian phase were obtained by creating a density plot.

### Induction of hyperthermic seizure

Following baseline recording, the mice were transferred to a glass-walled chamber equipped with a combined heating lamp and cooling mat. These experiments were conducted between 11:00 AM and 12:30 PM. Hyperthermia was induced using a heating lamp while cortical EEG and body temperature were continuously monitored. As soon as a seizure was detected, the mouse was placed on an iced mat to cool the body to 37°C, then returned to the recording cage where EEG monitoring was continued. The body cooling was used to improve clinical recovery from hyperthermia, however, SD incidence during hyperthermia in Scn1a^+/RX^ mice was also observed in initial studies without body cooling.

### Drug

Memantine was purchased from Tocris and PTZ was obtained from Sigma. Both drugs were dissolved in saline on the day of the experiment and administered by intraperitoneal injection.

### Study design and statistics

EEG recording studies included both male and female Scn1a^R1407X/+^ and Scn1a^+/+^ WT littermates in each cohort. Two to four mice were simultaneously recorded in each study cohort. Because some littermate mutant mice died or became moribund during the study, we added several Scn1a^RX^ mice, resulting in an unequal number of mice by sex. Cage location and order of treatments were randomly assigned and the experimenters were blinded to genotype when it is possible.

The number of animals in the initial characterization was determined based on power analysis. The numbers of animals in the memantine and PTZ studies were determined based on the results of the initial cohort.

Statistics were calculated by R and GraphPad prism software. Because most of the data showed skewed distribution, the statistical significances were tested using non-parametric methods such as aligned rank transformation ANOVA for multiple comparisons (77), Wilcox-signed rank test, and Mann-Whitney U-test for two sample comparisons. All two group comparisons were 2-tailed. A p-value less than 0.05 was considered significant. The data are presented as mean ± S.D unless otherwise mentioned in the figure legends.

### Study approval

All animal experiments were conducted under the guide of AAALAC and approved by IACUC of Baylor College of Medicine.

## Data availability

The raw data of this manuscript is included in the online supporting data. All electrophysiological and imaging data are available upon request

## Author Contributions

Conception and design of the study (IA), acquisition and analysis of data (IA, YN), manuscript draft preparation (IA, JLN). All authors contributed to the final draft of manuscript.

## Acknowledgments

This work was supported by an American Heart Association Career Development Grant 19CDA34660056 (to I.A.), the Curtis Hankamer Basic Research Fund at Baylor College of Medicine (I.A.), an American Epilepsy Society Junior Investigator Award (I.A.), the National Institutes of Health Center for SUDEP Research Grant NS090340 (to J.L.N.), and the Blue Bird Circle Foundation (J.L.N.).

